# A novel method for single whitefly (*Bemisia tabaci*) transcriptomes reveals an eleven amino acid deletion in the *NusG* protein in the bacterial endosymbiont *Portiera aleyrodidaru*

**DOI:** 10.1101/110353

**Authors:** P. Sseruwagi, J. M. Wainaina, J. Ndunguru, R. Tumuhimbise, F. Tairo, J. Guo, A. Vrielink, A. Blythe, T. Kinene, B. De Marchi, M.A. Kehoe, S.K. Tanz, L.M. Boykin

## Abstract

**Background:** *Bemisia tabaci* species (whiteflies) are the world’s most devastating insect pests within crops in the tropics. They cause billions of dollars (US) of damage each year and are leaving farmers in the developing world food insecure. Understanding the genetic and transcriptomic composition of these insect pests, the viruses they transmit and the microbiota is crucial to sustainable insect and virus management solutions for farmers. Currently, publically available transcriptome data for *B. tabaci* has been generated from pooled samples (mainly inbred lab colonies) consisting of several individuals because whiteflies are small (approximately 0.2 mm wide and 0.1 mm in height). Pooling individuals can lead to high heterozygosity and skewed representation of the genetic diversity. The ability to extract enough RNA from a single whitefly has remained elusive due to their small size and technology limitations. Therefore, the understanding of whitefly-microbiotad-viral species composition of an individual field-collected whitefly has also remained unknown. In this study, we developed a single whitefly RNA extraction procedure and subsequently successfully sequenced the transcriptome of four individual adult SubdSaharan Africa (SSA1) *B. tabaci.*

**Results:** Transcriptome sequencing on individual whiteflies resulted in between 39-42 million raw reads. *De novo* assembly of trimmed reads yielded between 65,000-162,000 transcripts across all four *B. tabaci* transcriptomes. In addition, Bayesian phylogenetic analysis of mitochondrion cytochrome I oxidase (mtCOI) grouped the four whiteflies within the SSA1 clade. BLAST searches on assembled transcripts within the four individual transcriptomes identified five endosymbionts; the primary endosymbiont *Portiera,aleyrodidarum* and four secondary endosymbionts: *Arsenophonus, Wolbachia, Rickettsia,* and *Cardinium spp.* These five endosymbionts were predominant across all four SSA1 *B. tabaci* study samples with prevalence levels of between 54.1d75%. Nucleotide and amino acid sequence alignments of the *Nus*G gene of *P. aleyrodidarum* for the SSA1 *B. tabaci* transcriptomes of samples WF2 and WF2b revealed an eleven amino acid residue deletion that was absent in samples WF1 and WF2a. Comparison of the protein structure of the *Nus*G protein from *P. aleyrodidarum* in SSA1 with known *Nus*G structures showed the deletion resulted in a shorter D loop. Although *Nus*G is key in regulating of transcription elongation, it is believed that the shortening of the loop region in the N-terminal domain is unlikely to affect transcription termination. Therefore, the effect of variability in this region across species is unknown.

**Conclusion:** In this study, we optimised a single whitefly high quality RNA extraction procedure and successfully carried out individual whitefly transcriptome sequencing on adult *B. tabaci* whiteflies. This enabled the detection of unique genetic differences in the *Nus*G genes of the primary endosymbiont *P. aleyrodidarum* in four field-collected SSA1 whiteflies that may not have been detected using lab-pooled *B. tabaci* isolines. The use of field-collected specimens means that both time and money will be saved in future studies using single whitefly transcriptomes in monitoring vector and viral interactions. In addition, the methods we have developed here are applicable to any small organism where RNA quantity has limited transcriptome studies.

## Background

Members of the whitefly *Bemisia tabaci* (Hemiptera: Aleyrodidae) species complex are classified as the world’s most devastating insect pests. There are 34 species globally [1] and the various species in the complex are morphologically identical. They transmit over 100 plant viruses [2, 3], become insecticide resistant [4], and ultimately cause billions of dollars in damage annually for farmers. The adult whiteflies are promiscuous feeders and will move between viral infected crops and native weeds that act as viral inoculum ‘sources’ and deposit viruses to alternative crops that act as viral ‘sinks’ while feeding.

The crop of importance for this study was cassava (*Manihot esculenta*). Cassava supports approximately 800 million people in over 105 countries as a source of food and nutritional security, especially within rural smallholder farming communities [5]. Cassava production in Sub Saharan Africa (SSA), especially the East Africa region is hampered by both DNA and RNA transmitted viruses.

Whitefly-transmitted viruses cause cassava mosaic disease (CMD) leading to 28-40% crop losses with estimated economic losses of up to $ 2.7 billion dollars per year in SSA [6]. The CMD pandemics in East Africa, and across other cassava producing areas in SSA were correlated with *B. tabaci* outbreaks [7]. African cassava mosaic viruses (ACMVs) occur mostly towards West Africa where a distinct group of *B. tabaci* SSA1 is predominant. On the other hand, East African cassava mosaic viruses (EACMVs) occur mainly in coastal areas of East Africa with highest diversity inland in Kenya, Tanzania and Uganda, yet again with a different group of *B. tabaci* SSA1. The two distinct groups of SSA1 are yet to be named. While some studies have been carried out to determine the relative transmission of CMDs by different *B. tabaci* species with indications of no significant differences, it is still not clear why some CMDs such as African cassava mosaic viruses (ACMVs) and East African cassava mosaic – Uganda variant (EACMVdUG), which is a recombinant between EACMV and ACMV in the coat protein (CP), do not occur in coastal East Africa.

Relevant to this study are two RNA *Potyviruses*: the Cassava Brown Streak viruses (CBSV) and the Uganda Cassava Brown Streak Virus (UCBSV) both devastating cassava in East Africa. *Bemisia tabaci*, species have been hypothesized to transmit these RNA viruses with limited transmission efficiency [8,9]. Recent studies have shown that there are multiple species of these viruses [10], which further strengthens the need to obtain data from individual whiteflies as pooled samples could contain different species with different virus composition and transmission efficiency. In addition, CBSV has been shown to have a higher rate of evolution than UCBSV [11] increasing the urgency of understanding the role played by the different whitefly species in the system.

### Endosymbionts and their role in B. tabaci

Viraldvector interactions within *B. tabaci* are further influenced by bacterial endosymbionts forming a tripartite interaction. *B. tabaci* has one of the highest numbers of endosymbiont bacterial infections with eight different vertically transmitted bacteria reported [12, 13], [14, 15]. They are classified into two categories; primary (P) and secondary (S) endosymbionts, many of which are in specialised cells called bacteriocytes, while a few are also found scattered throughout the whitefly body. A single obligate *P-symbiont P. aleyrodidarum* is systematically found in all *B. tabaci* individuals. *P. aleyrodidarum* is essential for whitefly survival as it supplies and complements the host metabolic activities in the synthesis of the essential amino acids threonine and tryptophan along with the non-essential amino acid serine [16]. *Portiera* has long co-evolutionary history with all members of the *Aleyrodinae* subfamily [16]. Although it is yet to be confirmed in whiteflies, most P-symbionts have been characteristically shown to have reduced and static genomes [17]. In this study, we further explore genes within the *P. aleyrodidarum* retrieved from individual whitefly transcriptomes, including the transcription termination/antitermination protein *Nus*G. *Nus*G is a highly conserved protein regulator that suppresses RNA polymerase pausing and increasing the elongation rate. However, its importance within gene regulation is species specific; in *Staphylococcus aureus* it is dispensable [18, 19].

The S-endosymbionts are not systematically associated with hosts and their contribution is not essential to the survival and reproduction. Seven facultative S-endosymbionts, *Wolbachia, Cardinium, Rickettsia, Arsenophonus, Hamiltonella defensa*, and *Fritschea, bemisae* have been detected in various *B. tabaci* populations [20, 21, 12, 22, 23]. The presence of S-endosymbionts can influence key biological parameters of the host. *Hamiltonella*, and *Rickettsia* facilitate plant virus transmission with increased acquisition and retention by whiteflies [24]. This is done by protection and safe transit of virions in haemolymph of insects through chaperonins (*GroEL*) and protein complexes that aid in protein folding and repair mechanisms [21].

### *Application of next generation sequencing in pest management of* B. tabaci

The advent of next generation sequencing (NGS) and specifically transcriptome sequencing has allowed the unmasking of this tripartite relationship of vector-viral-microbiota within insects [25, 26, 27]. Furthermore, NGS provides an opportunity to better understand the co-evolution of *B. tabaci* and its bacterial endosymbionts [28]. The endosymbionts have been implicated in influencing species complex formation in *B. tabaci* through conducting sweeps on the mitochondrial genome [29]. Applying transcriptome sequencing is essential to reveal the endosymbionts and their effects on the mitogenome of *B. tabaci* and predict potential hot spots for changes that are endosymbionts induced.

Several studies have explored the interaction between whitefly and endosymbionts and have resulted in the identification of candidate genes that maintain the relationship [30,31]. This has been explored as a source of potential RNAi pesticide control targets [32, 31, 32, 28]. RNAidbased pest control measures also provide opportunities to identify species-specific genes for target gene sequences for knock-down. However, to date all transcriptome sequencing has involved pooled samples obtained through rearing several generations of isolines of a single species to ensure high quantities of RNA for subsequent sequencing. This remains a major bottle neck in particular within arthropoda where collected samples are limited due to small morphological sizes [33, 34]. In addition, the development of isolines is time consuming and often has colonies dying off mainly due to inbreeding depression [35].

It is against this background that we sought to develop a method for single whitefly transcriptomes to understand the virus diversity within different whitefly species. We did not detect viral reads, probably an indication that the sampled whitefly was not carrying any viruses, but as proof of concept of the method, we validated the utility of the data generated by retrieving the microbiota *P-endosymbionts* and *S-endosymbionts* that have previously been characterised within *B. tabaci* [36, 37] In this study we report the endosymbionts present within fielddcollected individual African whiteflies and characterisation and evolution of the *Nus*G genes present within the *P-endosymbionts*.

## Results

### RNA extraction and NGS optimised for individual B. tabaci samples

In this study, we sampled four individual adult *B. tabaci* from cassava fields in Uganda (WF2) and Tanzania (WF1, WF2a, WF2b). Total RNA from single whitefly yielded high quality RNA with concentrations ranging from 69 ng to 244 ng that were used for library preparation and subsequent sequencing with Illumina Hiseq 2000 on a rapid run mode. The number of raw reads generated from each single whitefly ranged between 39,343,141 and 42,928,131 (Table 1). After trimming, the reads were assembled using Trinity resulting into 65,550 to 162,487 transcripts across the four SSA1 *B. tabaci* individuals (Table 1).

**Table 1.**
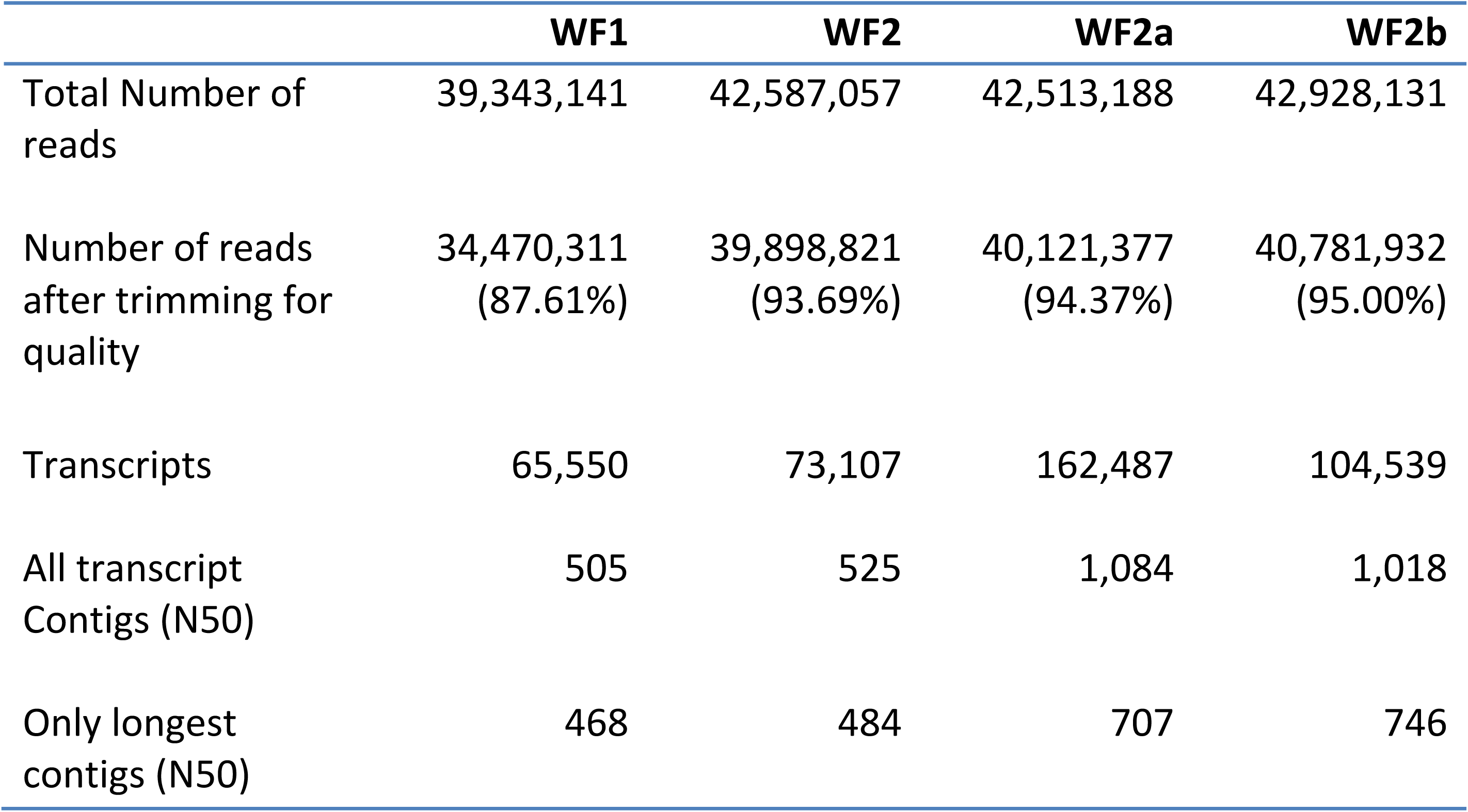
Summary Statistics from De novo trinity assemble of Illumina paired end individual whitefly transcriptome

### *Comparison of endosymbionts within the SSA1* B. tabaci *samples*

Comparison of the diversity of bacterial endosymbionts across individual whitefly transcripts was conducted with BLASTn searches on the non-redundant nucleotide database and by identifying the number of genes from each bacterial endosymbiont (Supplementary Table 1). We identified five main endosymbionts including: *P. aleyrodidarum* the primary endosymbionts and four secondary endosymbionts: *Arsenophonus, Wolbachia, Rickettsia sp, and Cardinium* spp (Table 2). *P. aleyrodidarum*, predominated all four SSA1 *B. tabaci* study samples with incidences of 74.8%, 71.2%, 54.1% and 58.5% for WF1, WF2, WF2a and WF2b, respectively. This was followed by *Arsenophonus, Wolbachia, Rickettsia sp*, and *Cardinium* spp, which occurred at an average of 18.0%, 5.9%, 1.6% and <1%, respectively across all four study samples.

**Table 2.**
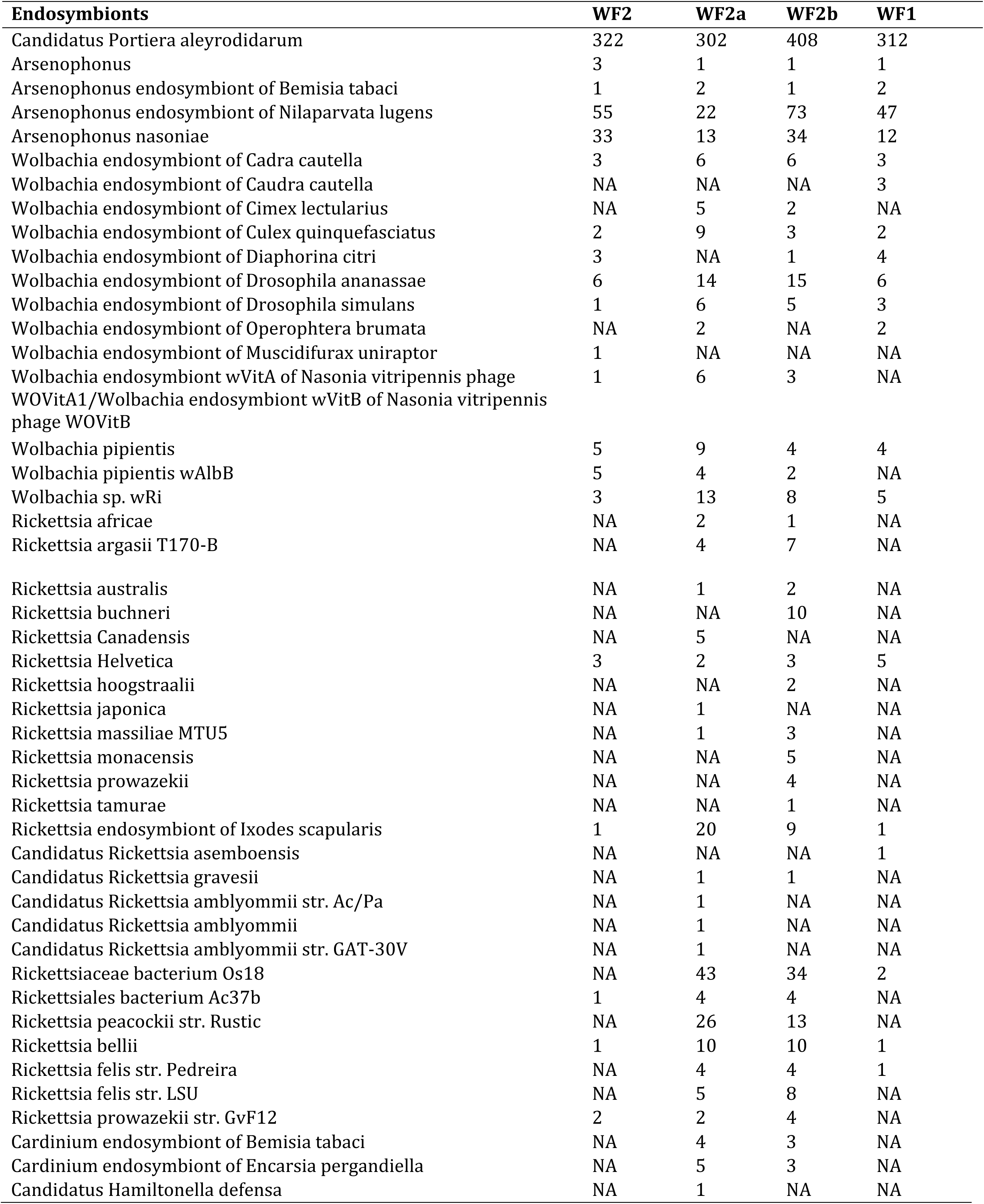
Distribution of endosymbionts and number of genes present in endosymbionts bacteria present in four SSA1 *B. tabaci* samples from this study

### Phylogenetic analysis of single whitefly mitochondrial cytochrome oxidase I (COI)

*B. tabaci* is recognized as a species complex of 34 species based on the mitochondrion cytochrome oxidase I [38, 1, 39]. We therefore used cytochrome oxidase I (COI) transcripts of the four individual whitefly to ascertain *B. tabaci* species status and their phylogenetic relation using reference *B. tabaci* COI GenBank sequences found at www.whiteflybase.org. All four COI sequences clustered within Sub Saharan Africa clade 1 (SSA1) species (data not shown).

### Sequence alignment and Bayesian phylogenetic analysis of NusG gene

Nucleotide and amino acid sequence alignments of the NusG in P. aleyrodidarum were conducted for several whitefly species including: *B. tabaci* (SSA1, Mediterranean (MED) and Middle East Asia Minor 1 (MEAM1) New World 2 (NW2), T. vaporariorum (Greenhouse whitefly) and *Alerodicus dispersus*. The alignment identified 11 missing amino acids in the *NusG* sequences for the SSA1 *B. tabaci* samples: WF2 and WF2b, *T. vaporariorum* (Greenhouse whitefly) and *Alerodicus disperses*. However, all 11 amino acids were present in samples WF1 and WF2a, MED, MEAM1 and NW2 (Fig. 1). Bayesian phylogenetic relationships of the *NusG* sequences of *P. aleyrodidarum* for the different whitefly species clustered all four SSA1 *B. tabaci* (WF1, WF2, WF2a and WF2b) within a single clade together with ancestral *B. tabaci* from GenBank (Fig. 2). The SSA1 clade was supported by posterior probabilities of 1 with *T. vaporariorum* and *Alerodicus*, which formed clades at the base of the phylogenetic tree (Fig. 2).

**Fig. 1.**
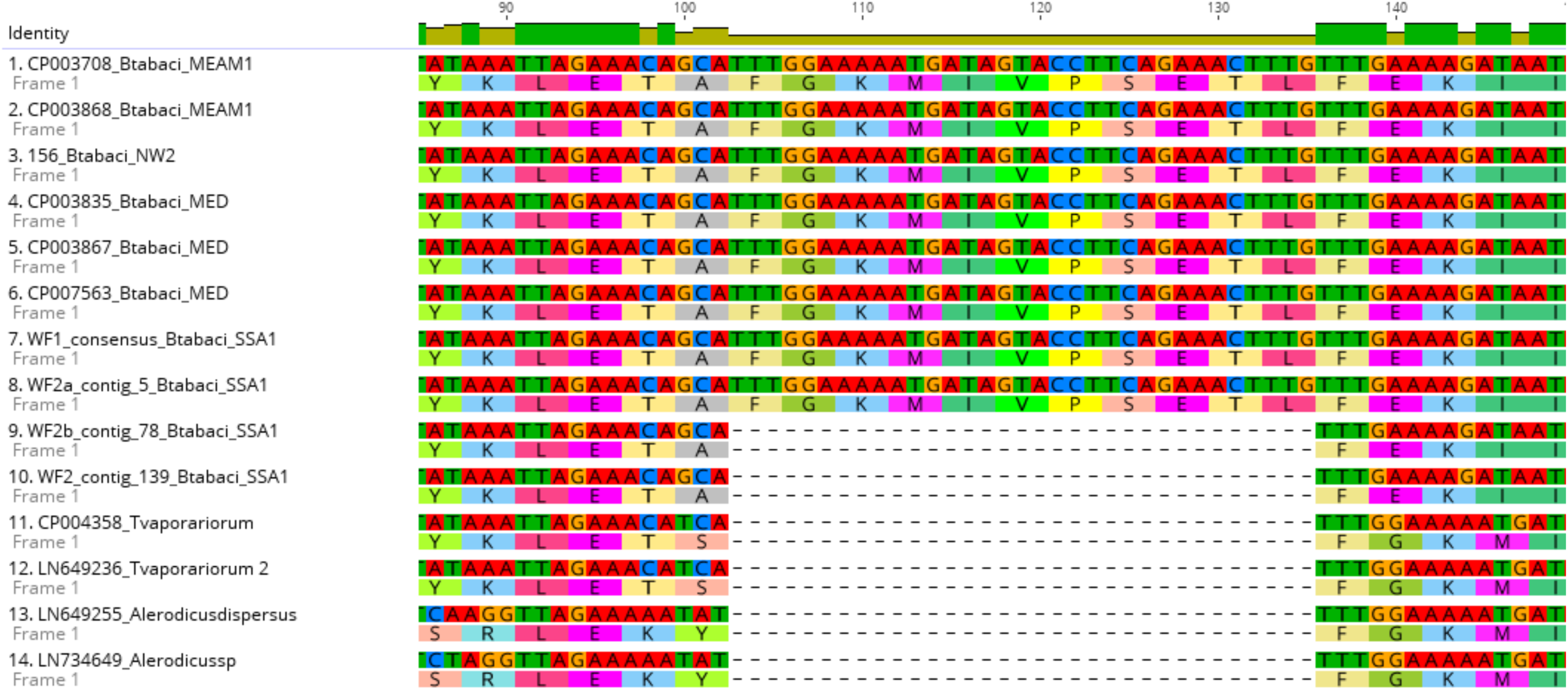
Sequence alignment of nucleotide sequences of *Nus*G gene in *P. aleyrodidarum* across whitefly species sequences using MAFFT v 7.017

**Fig. 2.**
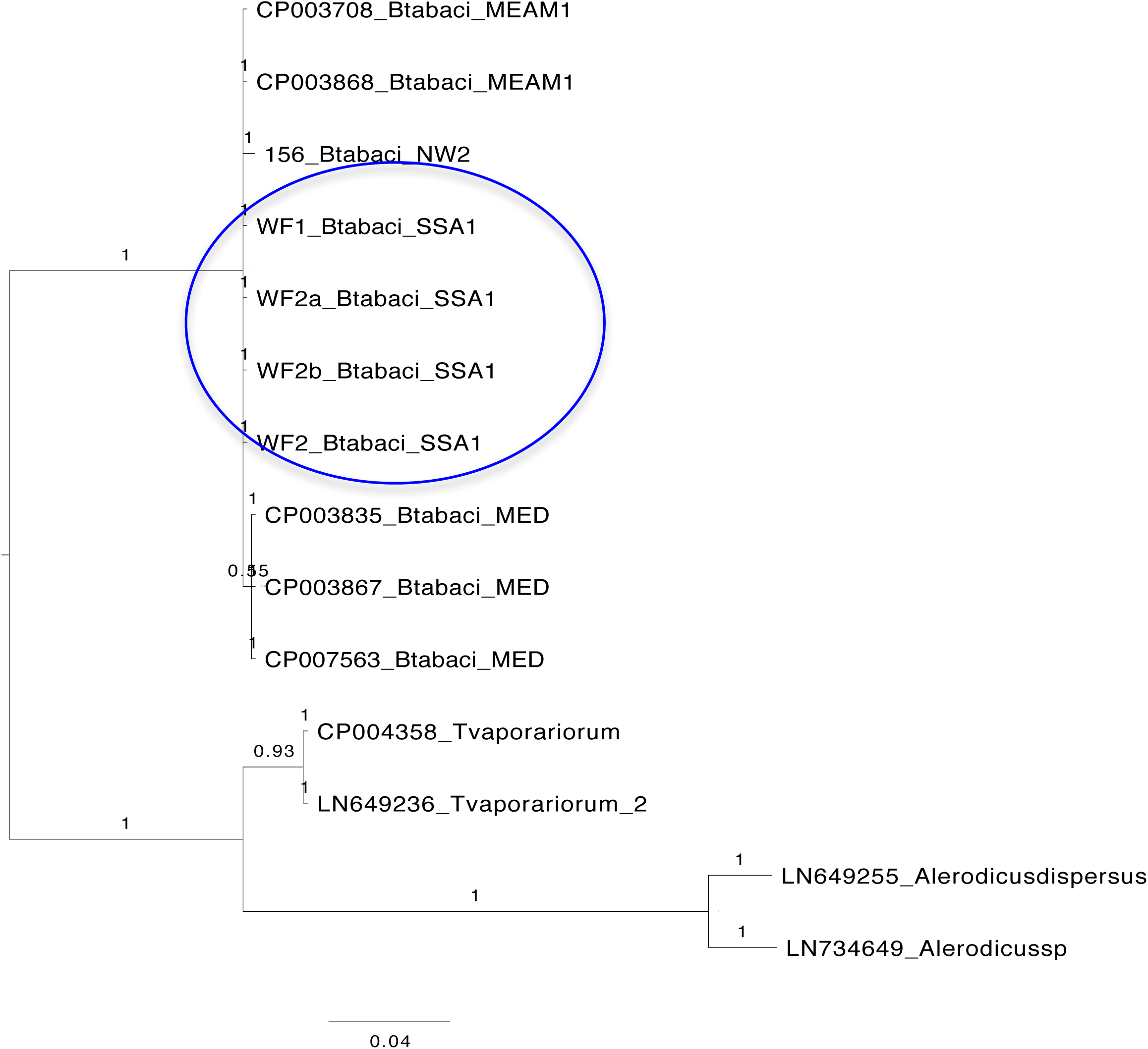
Bayesian phylogenetic tree of *Nus*G gene of *P. aleyrodidarum* across whitefly species using MrBayes -3.2.2 Circled are *B. tabaci* samples from this study

### Structure analysis of Portiera NusG genes

Structures of the *Nus*G protein sequence of the primary endosymbiont *P. aleyrodidarum* in the four SSA1 *B. tabaci* samples were predicated using Phyre2 with 100% confidence and compared to known structures of *Nus*G from other bacterial species including (*Escherichia, coli, Thermus thermophiles*, and *Aquifex aeolicus*; (PDB entries 2KO6, 1NZ8 and 1M1H, respectively) and Spt4/5 from yeast (*Saccharomyces cerevisiae;* PDB entry 2EXU) [18, 40, 41]. The 11-residue deletion was found in a loop region that is variable in length and structure across bacterial species, but is absent from archaeal and eukaryotic species (Fig. 3 and Fig. 4A). The effect of the deletion appears to shorten the loop in *Nus*G from the African whiteflies (WF2 and WF2b). A model of bacterial RNA polymerase (orange surface representation; PDB entry 2O5I) bound to the N-terminal domain of the T. thermophiles *Nus*G shows that the loop region is not involved in the interaction between *Nus*G and RNA polymerase (Fig. 4B).

**Fig. 3.**
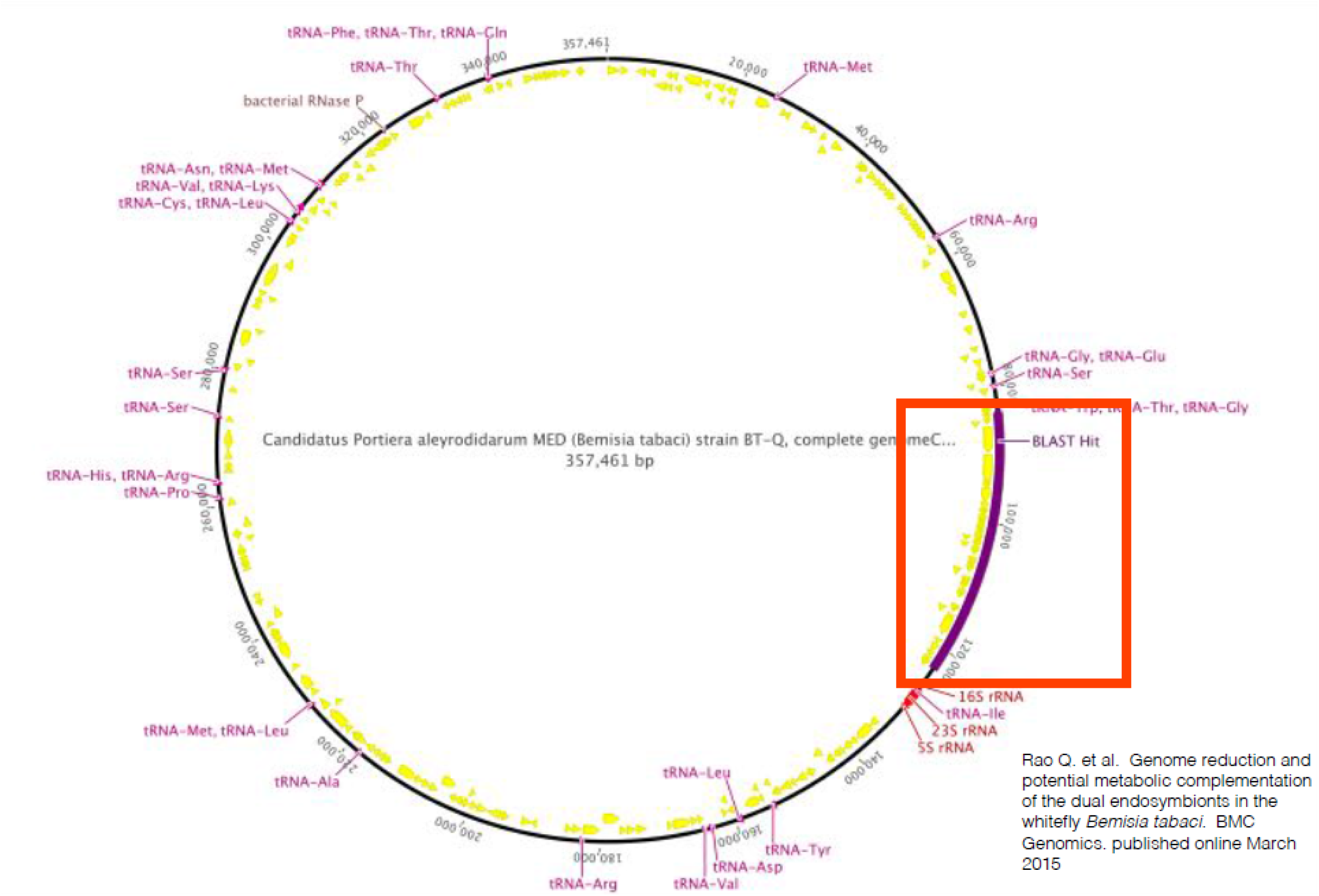
Structure of the *Nus*G gene showing the 11 amino acid deletion in a transcription factor of the primary endosymbiont *Portiera aleyrodidarum* of the SSA1 *B. tabaci* species

**Fig. 4.**
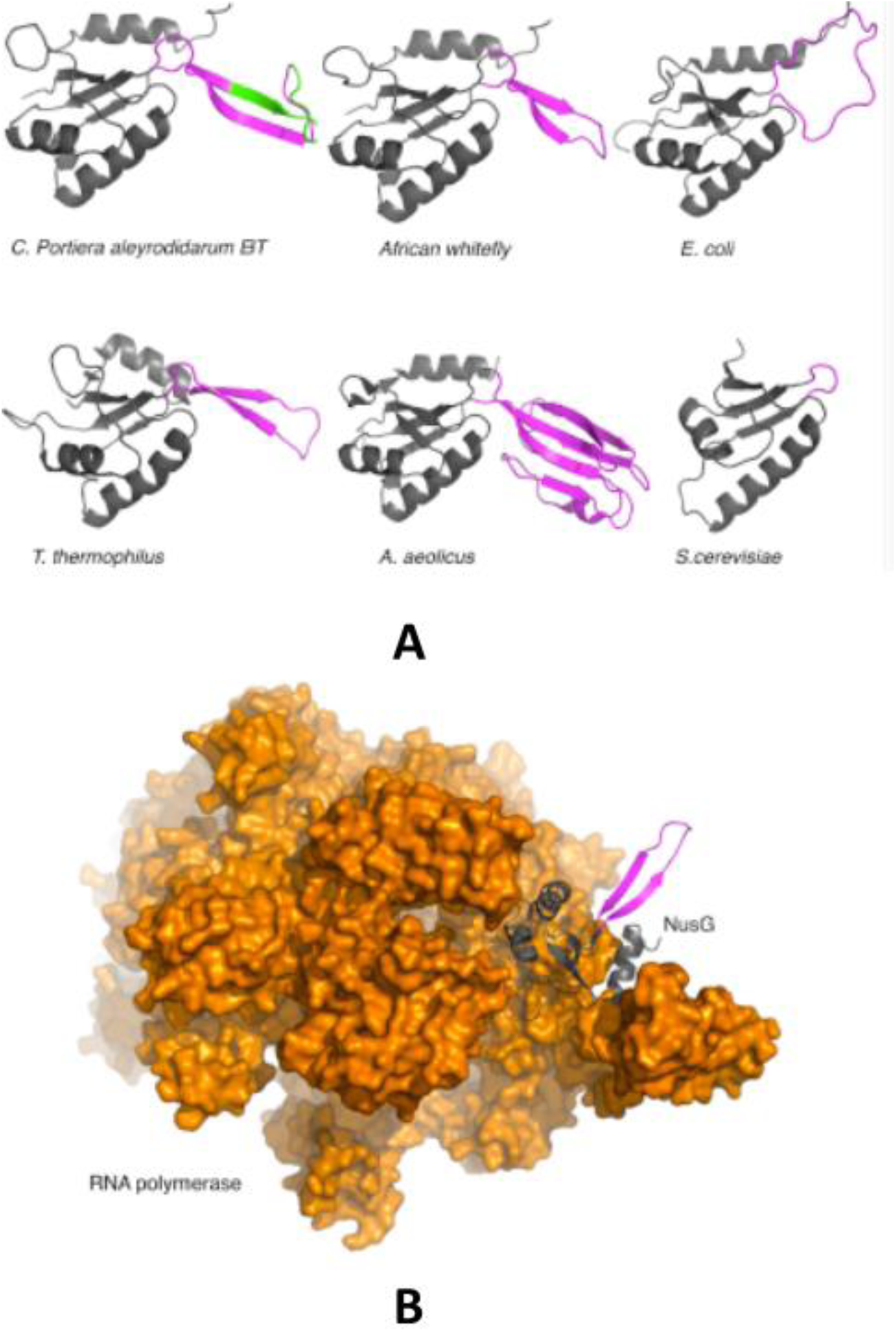
Structure analysis of *Nus*G from *P. aleyrodidarum* in *B. tabaci* and other endosymbionts **A.** Phyre2 based sstructure prediction of *Nus*G from *Candidatus Portiera aleyrodidarum* in *B. tabaci* SSAI whitefly and comparisons to the structures of *Nus*G from other bacterial species as indicated and of Spt4/5 from yeast. *Nus*G is coloured in grey, the loop region in magenta and the 11-residue deletion is shown in green in the C. *Portiera aleyrodidarum* structure. **B**. A model of bacterial RNA polymerase (orange surface representation) bound to the N-terminal domain of the *T. thermophiles Nus*G (grey cartoon representation)

## Discussion

In this study, we developed a single whitefly RNA extraction method for field-collected samples. We subsequently successfully conducted transcriptome sequencing on individual Sub-Saharan Africa 1 (SSA1) *B. tabaci*, revealing unique genetic diversity in the bacterial endosymbionts as proof of concept.

### *Nus*G deletion and implications within *P. aleyrodidarum* in SSA *B. tabaci*

We report the presence of the primary endosymbionts *P. aleyrodidarum* and several secondary endosymbionts within SSA1 transcriptome. Furthermore, *P. aleyrodidarum* in SSA1 *B. tabaci* was observed to have a deletion of 11 amino acids on the *Nus*G gene that is associated with cellular transcriptional processes within another bacteria species. On the other hand, *P. aleyrodidarum* from NW2, MED and SSA1 (WF2a, WF1) *B. tabaci* species did not have this deletion (Fig. 1). The deleted 11 amino acids were identified in a loop region of the N-terminal domain of *Nus*G protein resulting in a shortened loop in the SSA1 WF2b sample. This loop region has high variability in both structure and length across bacterial species and is absent from archaea and eukaryotic species.

*Nus*G is highly conserved and a major regulator of transcription elongation. It has been shown to directly interact with RNA polymerase to regulate transcriptional pausing and rho-dependent termination [19, 42, 18, 43]. Structural modelling of *Nus*G bound to RNA polymerase indicated that the shortened loop region seen in the WF2b sample is unlikely to affect this interaction. Rhoddependant termination has been attributed to the C-terminal (KOW) domain region of *Nus*G, therefore a shortening of the loop region in the N-terminal domain is also unlikely to affect transcription termination. Yet, there has been no function attributed to this loop region of *Nus*G, and thus the effect of variability in this region across species is unknown. However, the deletion could represent the result of evolutionary species divergence. Further sequencing of the gene is required across the *B. tabaci* species complex to gain further understanding of the diversity.

### Why the single whitefly transcriptome approach?

The sequencing of the whitefly transcriptome is crucial in understanding whitefly-microbiota-viral dynamics and thus circumventing the bottlenecks posed in sequencing the whitefly genome. The genome of whitefly is highly heterozygous [44]. Assembling of heterozygous genomes is complex due to the de Bruijn graph structures predominantly used [45]. To deal with the heterozygosity, previous studies have employed inbred lines obtained from raring a high number of whitefly isolines [46, 44, 33]. However, rearing whitefly isolines is time consuming and often colonies may suffer contaminations, leading to collapse and failure to raise the high numbers required for transcriptome sequencing.

We optimised the ARCTURUS^®^ PicoPure^®^ kit (Arcturus, CA, USA) protocol for individual whitefly RNA extraction with the dual aim of determining if we could obtain sufficient quantities of RNA from a single whitefly for transcriptome analysis and secondly, determine whether the optimised method would reveal whitefly microbiota as proof of concept. Using our method, the quantities of RNA obtained from field-collected single whitefly samples were sufficient for library preparation and subsequent transcriptome sequencing. Across all transcriptomes over 30M reads were obtained. The amount of transcripts were comparable to those reported in other arthropoda studies from field collections [34]. However, we did not observe any difference in assembly qualities as did [34]; probably due to the fact that our field-collected samples had degraded RNA based on RIN, and thus direct comparison with [34] was inappropriate.

Degraded insect specimen have been used successfully in previous studies [47]. This is significant, considering that a majority of insect specimen are usually collected under field conditions and stored in ethanol with different concentrations ranging from 70 to 100% [48, 49] rendering the samples liable to degradation. However, to ensure good keeping of insect specimen to be used for mRNA and total RNA isolation in molecular studies and other downstream applications such as histology and immunocytochemistry, it is advisable to collect the samples in an RNA stabilizing solution such as RNAlater. The solution stabilizes and protects cellular RNA in intact, unfrozen tissue and cell samples without jeopardizing the quality or quantity of RNA obtained after subsequent RNA isolation. The success of the method provided an opportunity to unmask vector-microbiota-viral dynamics in individual whiteflies in our study, and will be useful for similar studies on other small organisms.

### Endosymbionts diversity across individual SSA1 *B. tabaci* transcriptomes

In this study, we identified bacterial endosymbionts (Table 2) that were comparable to those previously reported in SSA1 *B. tabaci* on cassava [50, 23, 37]. Secondary endosymbionts have been implicated with different roles within *B. tabaci*. *Rickettsia* has been adversely reported across putative *B. tabaci* species, including the Eastern African region [51, 23, 51]. This endosymbiont has been associated with influencing thermo tolerance in *B. tabaci* species [52]. *Rickettsia* has also been associated with altering the reproductive system of *B. tabaci*, and within the females. This has been attributed to increasing fecundity, greater survival, host reproduction manipulation and the production of a higher proportion of daughters all of which increase the impact of virus [53]. *Arsenophonus, Wolbachia Arsenophonus* and *Cardinium spp* have been detected within MED and MEAM1 *Bemisia*species [12, 52]. In addition, [51] and [23] reported *Arsenophonus* within SSA1 *B. tabaci* in Eastern Africa that were collected on cassava. These endosymbionts have been associated with several deleterious functions within *B. tabaci* that include manipulating female-male host ratio through feminizing genetic males, coupled with male killing [54, 55].

Within the context of SSA agricultural systems, the role of endosymbionts in influencing *B.tabaci* viral transmission is important. Losses attributed to *B. tabaci* transmitted viruses within different crops are estimated to be in billions of US dollars [48]. Bacterial endosymbionts have been associated with influencing viral acquisition, transmission and retention, such as in *Tomato leaf curl virus* [56, 24]. Thus, better understanding of the diversity of the endosymbionts provides additional evidence on which members of *B. tabaci* species complex more proficiently transmit viruses and thus the need for concerted efforts towards the whitefly eradication.

## Conclusions

Our study provides a proof of concept that single whitefly RNA extraction and transcriptome sequencing is possible and the method is optimised and applicable to a range of small insect transcriptome studies. It is particularly useful in studies that wish to explore vector-microbiota-viral dynamics at individual insect level rather than pooling of insects. It is useful where genetic material is both limited as well of low quality, which is applicable to most agriculture field collections. In addition, the single whitefly transcriptome technique described in this study offers new opportunities to understand the biology and relative economic importance of the several whitefly species occurring in ecosystems within which food is produced in SubdSaharan Africa, and will enable the efficient development and deployment of sustainable pest and disease management strategies to ensure food security in the developing countries.

## Materials and methods

### Whitefly sample collection and study design,

In this study, we sampled whiteflies in Uganda and Tanzania from cassava (*Manihot esculenta*) fields. In Uganda, fresh adult whiteflies were collected from cassava fields at the National Crops Resources Research Institute (NaCRRI), Namulonge, Wakiso district, which located in central Uganda at 32°36′E and 0°31′N, and 1134 meters above sea level. On the other hand, the whiteflies obtained from Tanzania were collected on cassava in a countrywide survey conducted in 2013. The samples: WF2 (Uganda) and WF1, WF2a, and WF2b (Tanzania) used in this study were collected on CBSD-symptomatic cassava plants. In all the cases, the whitefly samples were kept in 70% ethanol in Eppendorf tubes until laboratory analysis. The whitefly samples were used for a two-fold function; firstly, to optimise a single whitefly RNA extraction protocol and secondly, to unmask RNA viruses and endosymbionts within *B. tabaci* as a proof of concept. In addition, data obtained from Nextera – DNA library prep from a Brazilian sample (156_NW2) was also used in this study. The whitefly was collected from a New World 2 colony in Brazil on *Euphorbia heterophylla* and kept in 70% ethanol in Eppendorf tubes until laboratory analysis.

### Extraction of total RNA from single whitefly

RNA extraction was carried out using the ARCTURUS^®^ PicoPure^®^ kit (Arcturus, CA, USA). Briefly, 30 µl of extraction buffer was added to an RNase-free micro centrifuge tube containing a single whitefly and ground using a sterile plastic pestle. To the cell extract an equal volume of 70% ethanol was added. To bind the RNA onto the column, the RNA purification columns were spun for two minutes at 100 x *g* and immediately followed by centrifugation at 16,000 x g for 30 seconds. The purification columns were then subjected to two washing steps using wash buffer 1 and 2 (ethyl alcohol). The purification column was transferred to a fresh RNAse-free 0.5 ml micro centrifuge tube, with 30 ul of RNAse-free water added to elute the RNA. The column was incubated at room temperature for five minutes, and subsequently spun for one minute at 1,000 x g, followed by 16,000 x g for one minute. The eluted RNA was returned into the column and re-extracted to increase the concentration. Extracted RNA was treated with DNase using the TURBO DNA free kit as described by the manufacturer (Ambion, life Technologies, CA USA). Concentration of RNA was done in a vacuum centrifuge (Eppendorf, Germany) at room temperature for 1 hour, the pellet was suspended in 15 ul of RNase-free water and stored at −80°C awaiting analysis. RNA was quantified, and the quality and integrity assessed using the 2100 Bioanalyzer (Agilent Technologies). Dilutions of up to x10 were made for each sample prior to analysis in the bioanalyzer.

### cDNA and illumina library preparation

Total RNA from each individual whitefly sample was used for cDNA library preparation using the Illumina TruSeq Stranded Total RNA Preparation kit as described by the manufacturer (Illumina, San Diego, CA, USA). Subsequently, sequencing was carried out using the HiSeq2000 on the rapid run mode generating 2 × 50 bp paired-end reads. Base calling, quality assessment and image analysis were conducted using the HiSeq control software v1.4.8 and Real Time Analysis v1.18.61 at the Australian Genome Research Facility (Perth, Australia).

## Analysis of NGS data using the supercomputer

### Assembly of RNA transcripts

Raw RNA-Seq reads were trimmed using Trimmomatic. The trimmed reads were used for *de novo* assembly using Trinity [57] with the following parameters: time -p srun --export=all -n 1 -c ${NUM_THREADS} Trinity --seqType fq --max_memory 30G --left 2_1.fastq --right 2_2.fastq --SS_lib_type RF --CPU ${NUM_THREADS} --trimmomatic --cleanup --min_contig_length 1000 -output _trinity min_glue = 1, V = 10, edgedthr = 0.05, min_kmer_cov = 2, path_reinforcement_distance = 150, and group pairs distance = 500.

### BLAST analysis of transcripts and annotation

BLAST searches of the transcripts under study were carried out on the NCBI (http://www.ncbi.nlm.nih.gov) non-redundant nucleotide database using the default cut-off on the Magnus Supercomputer at the Pawsey Supercomputer Centre Western Australia. Transcripts identical to known bacterial endosymbionts were identified and the number of genes from each identified endosymbiont bacteria determined.

### Phylogenetic analysis of whitefly mitochondrial cytochrome oxidase I (COI)

The phylogenetic relationship of mitochondrial cytochrome oxidase I (mtCOI) of the whitefly samples in this study were inferred using a Bayesian phylogenetic method implemented in MrBayes 3.2.2 [58]. The optimal substitution model was selected using Akaike Information Criteria (AIC) implemented in the jmodel test 2 [60].

### Sequence alignment and phylogenetic analysis of *Nus*G gene in *P. aleyrodidarum* across *B. tabaci* species

Sequence alignment of the *Nus*G gene from the P-endosymbiont *P. aleyrodidarum* from the SSA1 *B. tabaci* in this study was compared with another *B. tabaci* species, *Trialeurodes vaporariorum* and *Alerodicus dispersus* using MAFFT *v7.017* [61]. The Jmodel version 2 [60] was used to search for phylogenetic models with the Akaike information criterion selecting the optimal that was to be implemented in MrBayes 3.2.2. MrBayes run was carried out using the command: “lset nst=6 rates=gamma” for 50 million generations, with trees sampled every 1000 generations. In each of the runs, the first 25% (2,500) trees were discarded as burn in.

### *Analysis and modelling the structure of the Nus*G gene

The structures for *Portiera aleyrodidarum BT* and *B. tabaci* SSA1 whitefly were predicted using Phyre2 [62] with 100% confidence and compared to known structures of *Nus*G from other bacterial species. All models were prepared using Pymol (The PyMOL Molecular Graphics System, Version 1.5.0.4).

## Acknowledgements

J. M. W is supported by an Australian Award scholarship by the Department of Foreign Affairs and Trade (DFAT).

## Availabiliy of data

All raw reads for the four whitefly have been deposited in NCBI SRA under the accession SRR5110306, SRR5110307, SRR5109958

## Consent for publication

Not applicable

## Authors Contribution

LB, PS, JN, JMW ST designed the research the experiment. JMW, JG performed RNA-Seq analysis. RT, FR, BD, TK, MK provided samples and laboratory experiments. AV, AB did the NusG modelling. JMW, PS, LB wrote the manuscript. All authors read and approved the final manuscript.

